# Optimizing Motor Skills with HD-tDCS: Insights from a Pilot Study on Chopstick Proficiency

**DOI:** 10.64898/2026.01.30.702932

**Authors:** Jamie L. Scholl, Taylor J. Bosch, Lee A. Baugh

## Abstract

This study extends our previous research on neurological adaptations associated with learning to use chopsticks, in which we observed increased functional activity and connectivity changes in the anterior supramarginal gyrus (aSMG), a brain region previously implicated in novel tool use. In the present study, we investigated the effects of high-definition transcranial direct current stimulation (HD-tDCS) on motor learning by applying anodal stimulation to the aSMG in a double-blind, sham-controlled design with 24 participants (12 active, 12 sham). Participants in the active condition received ∼3 mA of HD-tDCS focused over the aSMG while watching a 20-minute video of the task – picking up a marble with chopsticks and dropping it into a cylindrical container. In comparison, participants in the sham condition watched the same video while receiving sham stimulation consisting of a 30-s ramp-up and ramp-down at the start and end of the 20-minute video. Immediately after the video task, all participants completed 15 one-minute trials in which they performed the task while electroencephalography (EEG) was recorded. Performance was assessed by the average number of successful marble drops per minute (MDPM) across trials. Additionally, video-based motion was analyzed using DeepLabCut to compare key kinematic metrics, providing insights into subtle variations in movement patterns during the marble task. Results showed a significant increase in MDPM in the active stimulation group compared to the sham group (17. 3 vs. 14. 1 MDPM; p < .05). Kinematic data showed increased movement jerk in the active stimulation group compared to sham (21719 vs 16926; p < .05), and EEG revealed differences in task-related gamma-band power over Cz (.0227 vs -.0758; p < .05). These findings suggest that HD-tDCS enhances the rate of motor learning in novel tool use and underscore the potential of aSMG-targeted stimulation in facilitating complex motor tasks. Further studies are warranted to explore the broader applicability of HD-tDCS in skill acquisition and rehabilitation.

**New and Noteworthy:** The presented study shows the role that the left anterior supramarginal gyrus plays in experience-independent tool learning. Anodal HD-tDCS applied during action observation increased performance in a subsequent chopstick skill acquisition task. This increase in performance was accompanied by enhanced task-related gamma-band activity and altered movement kinematics. By linking neuromodulation of the parietal tool-use hub to behavioral, kinematics, and electrophysiological changes, these findings significantly advance our understanding of how higher-order sensorimotor networks support tool-use learning.

## Introduction

The use of novel tools is crucial in humans as it showcases our advanced ability to integrate motor planning and action [1]. This skill allows humans to visualize a sequence of steps before executing complex, precise movements. Tool use also involves learning, adaptability, and problem-solving, as well as the ability to adjust actions based on feedback. This coordination between planning and motor control is fundamental in tasks ranging from daily activities to innovations, emphasizing the role of tool use in human evolution and cognitive development [1, 2].

Neuroimaging and comparative neuroscience research have identified a unique neural structure in humans, the left anterior supramarginal gyrus (aSMG). This region has been consistently implicated in tool use and tool-related action observation, showing robust activation when individuals observe others using tools, even when the mode of tool use varies [3-6]. Importantly, activation of the aSMG during action observation appears to be independent of direct motor execution, suggesting a role in higher-order representations of tool-related actions rather than simple motor mirroring.

Recent work further suggests that the left aSMG functions as an integrative hub within the human tool-use network. Specifically, it is thought to combine sensorimotor information from the anterior intraparietal sulcus (aIPS) with semantic knowledge and technical reasoning processes. When observing unfamiliar tool use, the left aSMG merges semantic information with technical reasoning, enabling the selection of the appropriate object affordance and planning of a grasping action. This suggests that the left aSMG acts as a key hub in a unique human tool-use network, facilitating experience-independent representations of unfamiliar tools [7]. This is further supported by neuromodulation work to transiently interfere with the left SMG in healthy participants performing judgment tasks about hand configuration in to the context of tool use. Following inhibitory stimulation of the SMG, participants demonstrated impaired hand configurations independent of contextual judgments, further emphasizing the role of this region in the processing of incoming information about tools to facilitate the shaping of hand posture [8].

Building on this framework, the present study examined whether modulating activity in the left aSMG with high-definition transcranial direct current stimulation (HD-tDCS) during action observation enhances subsequent motor learning of a previously unfamiliar tool (chopsticks). Chopsticks have previously been used to examine the motor learning process [9] and provide a role for technical reasoning in the tool-use network. This study proposes that increasing neural activity in the left anterior supramarginal gyrus (aSMG) via transcranial direct current stimulation during the observation of novel tool use, prior to any behavioral experience, will enhance the rate of motor learning. If the hypotheses regarding the left aSMG are validated, it would provide additional evidence that this area is key in facilitating experience-independent tool use in humans. This finding is particularly significant given the rising prevalence of virtual tool use in everyday life, which is believed to rely on the same underlying neural networks [10].

In conjunction with neuromodulation, video recordings will enable quantitative kinematic analysis. In the context of tool use, these measures are particularly informative because skilled performance often emerges through gradual refinement of movement structure rather than immediate task success. Prior work has demonstrated that kinematic signatures can reliably differentiate novice from expert tool users, reveal learning-related changes across practice, and identify compensatory strategies following neural disruption [11-13]. Video-based kinematic approaches have also been instrumental in examining how individuals acquire novel tool skills, such as chopstick use, by capturing changes in grasp configuration, movement variability, and coordination patterns over time [9]. Markerless video tracking methods have further expanded the utility of kinematic analysis by enabling high-resolution, naturalistic assessment of tool-related actions without constraining movement [14]. Together, kinematic video-based analyses provide insight into the underlying mechanisms of motor learning and sensorimotor integration, making them particularly well-suited for examining changes in acquisition and optimization of novel tool use following neuromodulation.

We were also interested in electroencephalographic changes that coincided with stimulation versus sham stimulation. Specifically, we were interested in examining both beta (∼13-30 Hz) and gamma (>30 Hz) event-related oscillatory activity because these frequencies have been shown to be correlated with control of voluntary movement and motor learning. Beta oscillations within the sensorimotor network are known to be suppressed prior to and during movement (event-related desynchronization) and also show a post-movement rebound. This pattern is thought to reflect the release and subsequent re-stabilization of the motor system during action and sensorimotor updating [15, 16]. It has also been shown that beta band dynamics are sensitive to practice and learning, with observed changes in beta power and timing being linked to the acquisition and refinement of motor skills and to the updating of internal models supporting skilled action [17, 18]. In comparison to beta-band activity, gamma band activity in motor cortical regions is known to increase around movement onset and during movement execution, with this activity being though to reflect localized cortical processing related to movement kinematics, force control, and the integration of ongoing sensory feedback [19-21]. Importantly, gamma oscillations have been particularly associated with precision grasping and fine motor control, skills that are critical for skilled tool use [20]. Since skilled chopstick manipulation requires continuous adjustments in grip force, coordination, and online error correction, measuring event-related spectral perturbations in the beta and gamma bands may provide a window into the cortical processing underlying the motor learning task.

Building on this prior work, the present study tested the hypothesis that anodal HD-tDCS applied during observational learning would increase subsequent behavioral performance in a chopstick-based marble manipulation task, as evidenced by an increase in the number of successful marble drops per minute relative to sham stimulation. We further hypothesized that stimulation-related performance gains would be reflected in measurable changes in movement kinematics, reflecting more efficient motor execution, including increased movement speed and altered total path length during task performance. At the neural level, we hypothesized that enhanced motor learning following active stimulation would be accompanied by modulation of sensorimotor cortical dynamics, particularly in beta- and gamma-band power at central electrode sites (C3, Cz, C4). Together, these hypotheses were designed to test whether targeted neuromodulation of the aSMG enhances motor learning by coordinating behavioral, kinematic, and electrophysiological mechanisms.

## Materials and Methods

### Participants

Thirty-three participants were recruited from the University of South Dakota (17 female, 15 male, 1 nonbinary; mean age = 19.19 years; age range = 18-23 years). All participants provided written informed consent in accordance with the experimental procedures approved by the University of South Dakota Institutional Review Board and the Declaration of Helsinki.

Participants were screened for eligibility and their ability to receive neurostimulation safely (e.g., no neurological or psychological conditions that are contraindicated for tDCS), were right-hand dominant, and had minimal prior experience using chopsticks was assessed using a subjective questionnaire requiring participants to rank their ability to skillfully use chopsticks (scale = 1-6; inclusion criterion was a ranking ≤ 3). Additionally, the screening asked for a self-report of chopstick use frequency (never to daily), with once/month or more resulting in exclusion.

Electroencephalography (EEG) was utilized to record neural activity throughout the stimulation and behavioral trials. Data was collected via a 64-channel passive EEG system with the extended International 10-10 montage (g.Hlamp-Research with g.LADYBIRD electrode, g.tec Medical Engineering, Austria). Signals were acquired at 1000 Hz, and filtered online to remove nonphysiological signals. EEG recordings were collected across the stimulation timeframe (20 minutes) as well as throughout the behavioral testing. Collected data was manually inspected for artifacts, and analyses followed standard preprocessing steps. Raw continuous EEG recordings were imported into EEGLAB using a custom MATLAB script. Following import, all datasets were downsampled to 300 Hz to reduce computational load while preserving frequencies of interest for subsequent analyses. Continuous EEG data were band-pass filtered using EEGLAB’s basic finite impulse response (FIR) filter. A high-pass filter at 1 Hz was applied to remove slow drifts and low-frequency noise. A low-pass filter at 100 Hz was applied to attenuate high-frequency noise while retaining beta-band and gamma-band activity. Noisy or malfunctioning channels were identified using automated and visual inspection procedures. Channels were flagged based on criteria including flatline detection, excessive noise, abnormal probability distributions, or kurtosis. Flagged channels were visually inspected using scroll plots of channel time series and subsequently removed prior to re-referencing and independent component analysis (ICA). After channel rejection, EEG data were re-referenced to the common average reference. Extended Infomax ICA was performed on the continuous, filtered, and average-referenced data to separate neural and non-neural signal sources. All retained EEG channels were included in the ICA decomposition (AMICA). Resulting ICA weights were saved for subsequent component classification and rejection. Independent components were classified using the ICLabel algorithm. Components were evaluated based on their scalp topographies, time courses, and spectral properties. Automatic component rejection was performed using ICLabel probability thresholds of ≥ 0.80 for non-brain classes, including eye, muscle, cardiac, line noise, channel noise, and “other” components. Components exceeding these thresholds were marked and removed from the data. Cleaned datasets were saved following component rejection. Cleaned continuous EEG data were segmented into epochs time-locked to task-related events. Epochs extended from −2.0 s to +60.0 s relative to event onset. Following epoching, noisy trials were optionally rejected using automated epoch rejection methods based on joint probability, kurtosis, or amplitude thresholds. Only clean, artifact-free epochs were retained for further analyses.

Time–frequency analysis was performed using EEGLAB’s pop_newtimef function to compute event-related spectral perturbations (ERSPs) on a single-trial basis. ERSPs were computed separately for three sensorimotor electrodes of interest: C3, Cz, and C4. For each electrode, ERSPs were calculated using Morlet wavelet convolution with a wavelet factor of 4 cycles and a linear increase in cycles (0.5 cycles per frequency). Power estimates were computed over a time window extending from −2000 ms to +60,000 ms relative to epoch onset, with 200 output time points evenly spaced across the epoch. A pre-event baseline from −2000 to 0 ms was used for baseline correction. Spectral power was estimated within two a priori frequency bands of interest (Beta band: 15–30 Hz; Gamma band: 30-100 Hz). For each frequency band and electrode, ERSP values were retained as a three-dimensional matrix (frequency × time × trial). Complex values returned by the time–frequency decomposition were converted to real-valued power estimates prior to further analysis. For each trial, mean spectral power was computed by averaging ERSP values across all frequencies and time points within the specified frequency band. This procedure yielded one scalar power value per trial for each electrode–frequency band combination (C3 beta, C3 gamma, Cz beta, Cz gamma, C4 beta, and C4 gamma). These values were stored as trial-level measures for subsequent statistical analysis [22].

Participants were randomly assigned to receive either ∼3mA of HD-tDCS stimulation to the left aSMG or sham stimulation prior to the 15 behavioral sessions. Both the participant and the experimenter were unaware of the experimental condition, with only the stimulation machine operator being aware of condition. The remaining experimental team members remained blinded until all data collection was complete. Stimulation was delivered using a 4x1 HD-tDCS stimulator (Soterix Medical Inc, New York, NY) via Ag/AgCl sintered ring electrodes. Parameters and current modeling [23] were designed to deliver maximum stimulation to the aSMG (Figure 1). Participants assigned [23]d to receive sham stimulation had electrodes placed in an identical montage as the active group. However, the current was only applied for 60 seconds (30-second ramp-up, 30-second ramp-down) at the start and end of the stimulation protocol. This process has previously been reported as a reliable method for blinding participants to their assignment in tDCS trials, with no substantial effect on cortical excitability [24]. Stimulation (active or sham) was applied for 20 minutes, during which participants watched a video of the action they were expected to learn. These videos depicted an actor skillfully using chopsticks to perform the behavioral task from a first-person perspective and were presented using custom LabVIEW software (LabVIEW 2014, National Instruments). Specifically, experimental stimuli included a video depicting an actor grasping a marble (diameter of marble = 16mm) with a pair of bamboo chopsticks (length of chopsticks = 260mm), lifting the marble to the apex of a cylindrical container (height of cylinder = 195mm), and dropping the marble into the cylindrical container (diameter of cylinder = 45mm).

**Figure 1.**
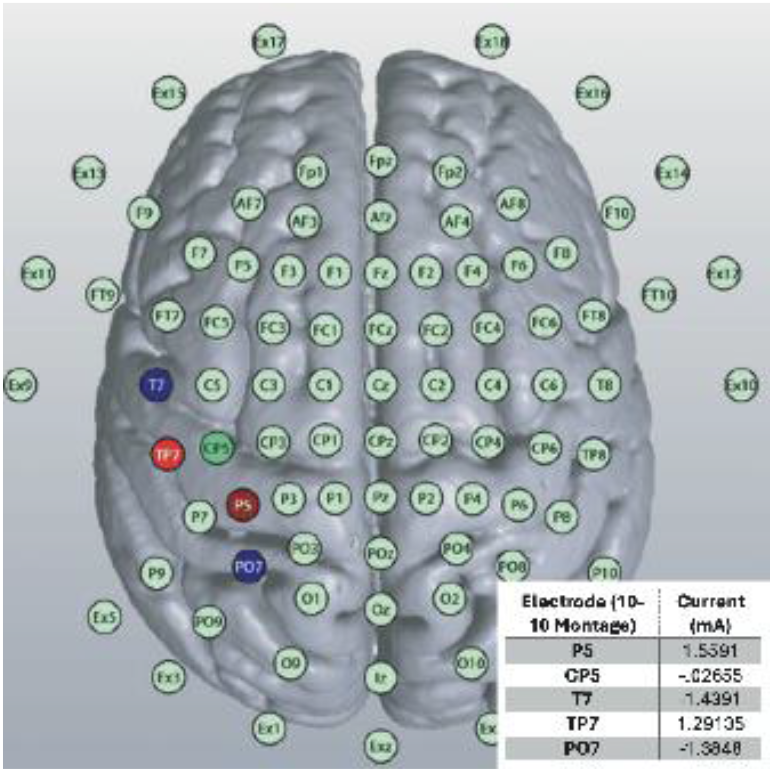
Current stimulation model for the left aSMG at ∼3mA. The electrode site, current, and field intensity are displayed following modeling on the MNI152. L=Left.

Immediately following the stimulation protocol, the participants began the behavioral task. Each behavioral training session consisted of 15 trials (duration = 60 seconds per trial), during which each participant used chopsticks to grasp a marble and drop it into the cylindrical container as many times as possible over the trial’s duration (Figure 2). Participants used practice chopsticks to perform the task, which included a loop for both the index and middle fingers, as well as a bridge connecting the two chopsticks. Using these practice chopsticks in each session ensured that all participants learned to use them consistently, while minimizing the overall difficulty of the task. The number of times participants successfully dropped the marble into the cylindrical container was recorded manually by the experimenter during each trial and later analyzed from video recordings.

**Figure 2.**
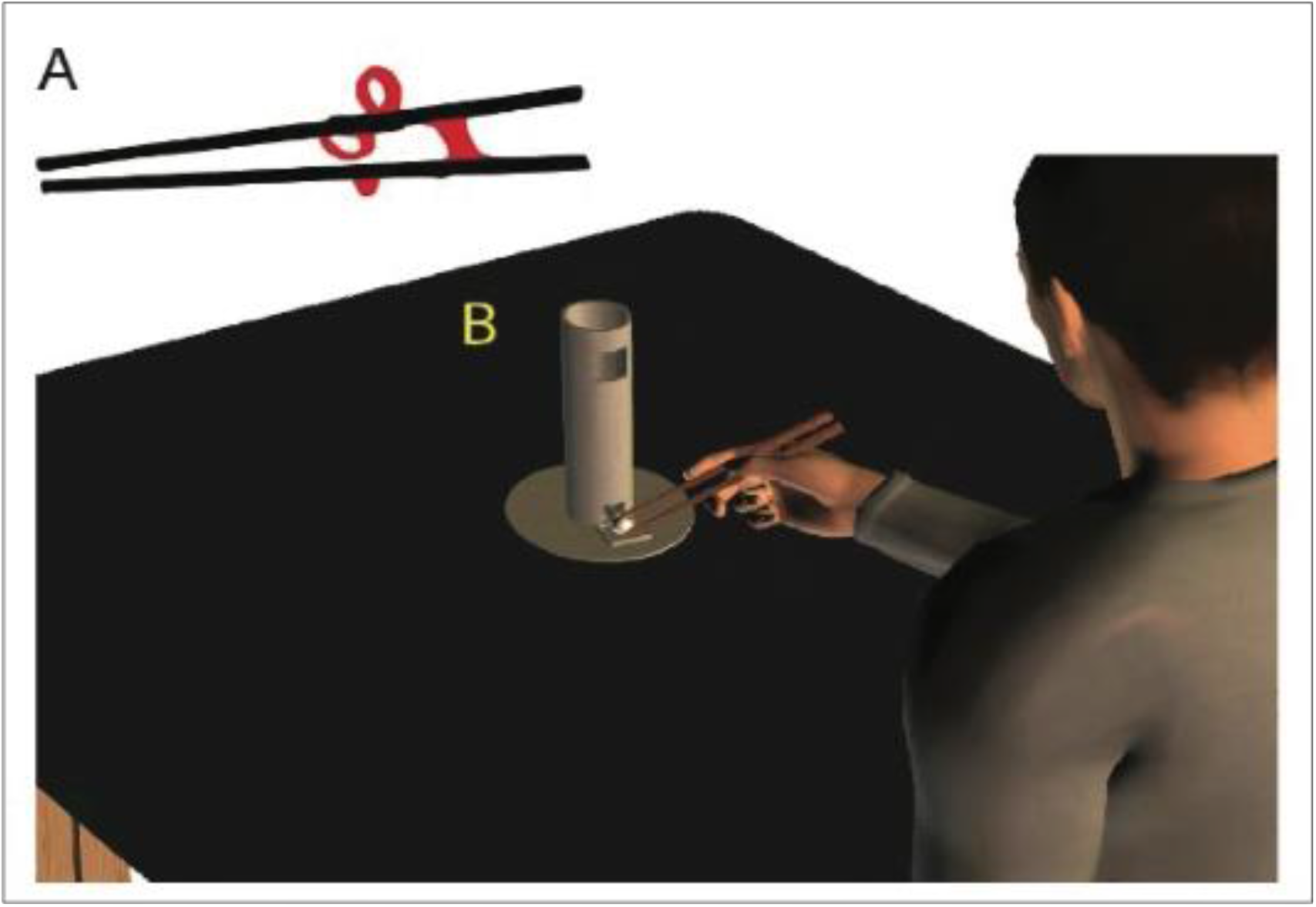
Practice chopsticks were used to ensure all participants learned to hold the chopsticks in a similar manner (A). Participants were instructed to pick up the marble with the chopsticks held in their dominant hand (right) and place it in the top of the cylinder as many times as possible during each trial (B).

Video recordings were made using a Basler acA1300-75gc camera consisting of a 1.3MP machine vision camera recording with a 1280 x 1024 pixel resolution and 30 Hz frame rate from a fixed location tripod. Camera control and capture was performed by custom LabVIEW software. Frames for labeling were extracted uniformly across each video, and all labeled data were normalized prior to training. Markerless pose estimation was performed using DeepLabCut with a PyTorch backend. Nine landmarks (See Figure 3) were manually labeled, including anatomical landmarks of the proximal interphalangeal joint and the distal phalanx of the second and third phalanges, the metacarpophalangeal joint and distal phalanx of the first digit, and the center of the wrist, over the scaphoid bone, all on the right side. Other landmarks included the tip of the top and bottom chopstick, and the LED light located at the top of the cylindrical container. Pose estimation was performed using a deep convolutional neural network with a ResNet-50 backbone incorporating group normalization (ResNet-50-GN) and an output stride of 16. The network used a heatmap-based keypoint detection head with integrated location refinement. For each body part, a Gaussian heatmap representation was generated, with simultaneous prediction of subpixel offsets using a location refinement branch. Heatmap prediction loss was computed using a weighted mean squared error criterion, while location refinement loss was computed using a weighted Huber loss. Extensive data augmentation was applied during training to improve generalization. Augmentations included random affine transformations (rotation up to ±30°, scaling between 0.5× and 1.25×, applied with 50% probability), hybrid crop sampling with a target crop size of 448 × 448 pixels and a maximum spatial shift of 10%, additive Gaussian noise, and motion blur. Input images were normalized prior to inference and training. The network was trained using the AdamW optimizer with an initial learning rate of 5 × 10^−4^. A staged learning rate schedule was employed, with learning rate reductions at epochs 90 and 120. Training was performed for a total of 200 epochs using a batch size of 8. Model evaluation was performed every 10 epochs, and up to 5 model snapshots were saved every 25 epochs. Random initialization was controlled using a fixed seed. Model performance was evaluated on a held-out test set using DeepLabCut’s standard evaluation pipeline. Evaluation was performed using a likelihood cutoff (pcutoff) of 0.6. For the final selected model, the mean root mean squared error (RMSE) on the training set was 6.97 pixels, which decreased to 2.91 pixels when excluding low-confidence predictions below the likelihood cutoff. Correspondingly, training performance achieved a mean average precision (mAP) of 96.81% and a mean average recall (mAR) of 98.08%. On the held-out test set, the final model achieved an RMSE of 6.62 pixels, which improved to 2.83 pixels after applying the likelihood cutoff. Test set performance reached a mean average precision (mAP) of 98.86% and a mean average recall (mAR) of 99.26%, indicating strong generalization across body parts and minimal overfitting. The final model snapshot was selected based on maximal test set mAP.

**Figure 3.**
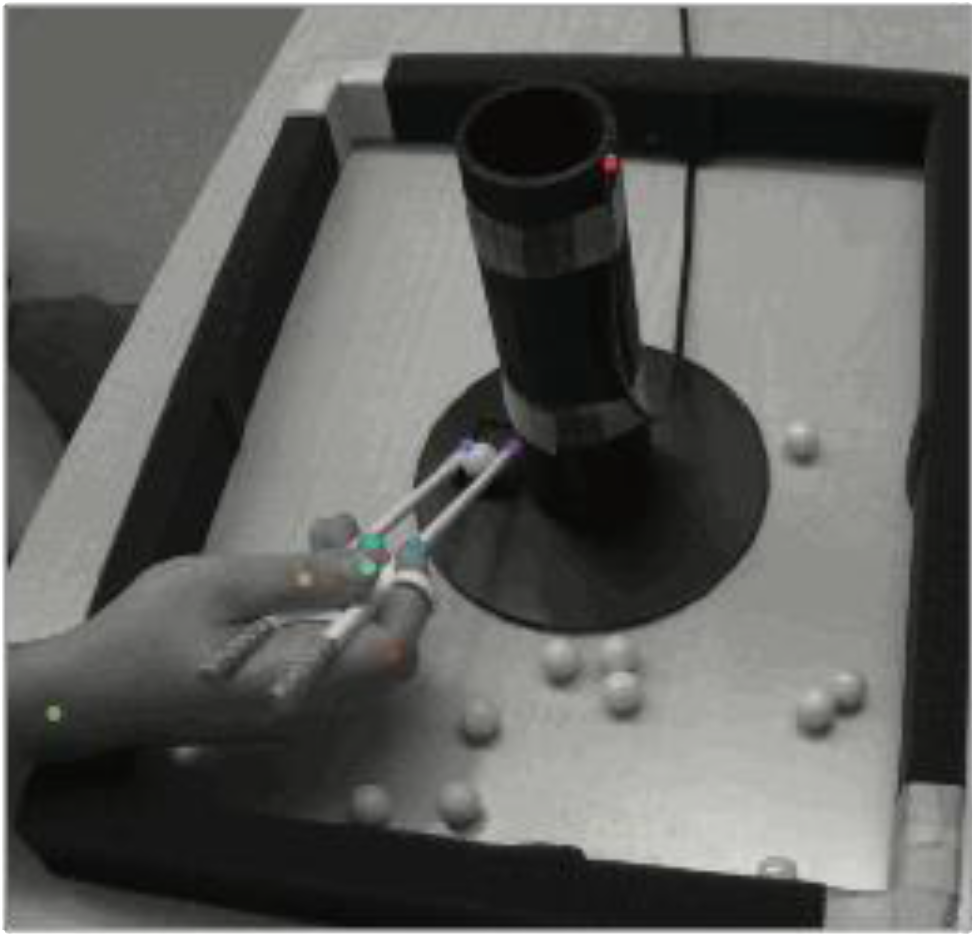
Example of the nine landmark locations used for markerless pose estimation within DeepLabCut with a PyTorch backend. Landmarks included the proximal interphalangeal joint and the distal phalanx of the second and third phalanges, the metacarpophalangeal joint and distal phalanx of the first digit, and the center of the wrist, over the scaphoid bone, all on the right side. Additionally, the tips of the top and bottom chopstick and the LED light located at the top of the cylindrical container were labeled.

Statistical analyses were conducted using jamovi [22], which operates on the R statistical environment (R version 4.5). All statistical tests were two-tailed with an alpha level of 0.05.

Behavioral results were quantified as the number of successful marble drops per minute (MDPM) during each of the fifteen trials. To evaluate differences in task performance between stimulation conditions (active stimulation vs. sham), a between-subjects one-way analysis of variance (ANOVA) was performed on the average behavioral outcome measure (marble drops per minute). Because group variances were not assumed to be equal, Welch’s ANOVA was used, with adjusted degrees of freedom reported. Kinematic data were analyzed for total path length (px) and movement velocity (px/s) of the top chopstick tip. In addition to conventional kinematic measures of movement performance, we quantified movement jerk, defined as the first temporal derivative of acceleration. Jerk is a well-established metric of movement smoothness and motor control stability, with higher jerk values reflecting more abrupt changes in force production and less smooth motor execution [25, 26]. Thus, jerk was included to capture potential stimulation-induced alterations in motor planning, online correction, and control strategy that may not be reflected by speed-based measures alone. Lastly, EEG data from C3, Cz, and C4 were compared using independent samples t-tests, also with adjusted degrees of freedom.

## Results

All participants were screened for familiarity with chopstick use, resulting in the exclusion of 5 participants’ data from the final dataset. The final mean familiarity for chopstick use (reported as M ± SEM) was 1.39 ± 0.10 on a six-point scale, validating that the participants were naïve to the use of chopsticks prior to training. An additional 2 participants were excluded for reporting the use of chopsticks more than once per month on their screening form. Lastly, two participants were removed for reporting chopstick use of once per year, but with initial trial scores more than 3.5 standard deviations above the mean of all participants, representative of proficient chopstick abilities at the start of the experiment. This resulted in a final participant count of 24 participants (12 stimulation, 12 sham stimulation; male = 12, female = 11, nonbinary = 1), with a mean age of 19.4 years (range: 18–23). One additional sham participant was removed from the kinematic data due to equipment malfunction preventing the recording of the kinematic videos.

For MDPM, we performed a one-way ANOVA, which revealed a significant main effect of stimulation on behavioral performance (Figure 4; F(1,21.4) = 5.80; p = 0.025). Participants receiving active stimulation exhibited a higher average rate of marble drops per minute compared to the sham group (17.3 MDPM vs. 14.1 MDPM). Kinematic data measured by total path length and speed were plotted over still images (Figure 5). Compared with the sham control group, participants receiving active stimulation had an increased movement jerk (Figure 6A; 21719 vs 16926; t(20.4) = 2.819, p = .010. Neither path length (Figure 6B; 10284 pixels vs. 88299 pixels; t(16.5) = 1.52; p = 0.147) nor movement velocity (Figure 6C; 1909 pps vs. 1626 pps; t(15.8) = 1.8; p = 0.073) were significantly different between active and sham stimulation. Condition-related differences in task-related gamma-band power were observed over Cz, with higher gamma power during active stimulation (.0227) compared with to sham stimulation (-.0758) (Figure 7; (t(14.2) = 3.58, p = .003). No significant differences were observed at C3 (p >.10) or at C4 (t(21.9) = 2.07, p = .051). Additionally, there were no significant differences found when examining beta power (p > 0.10).

**Figure 4.**
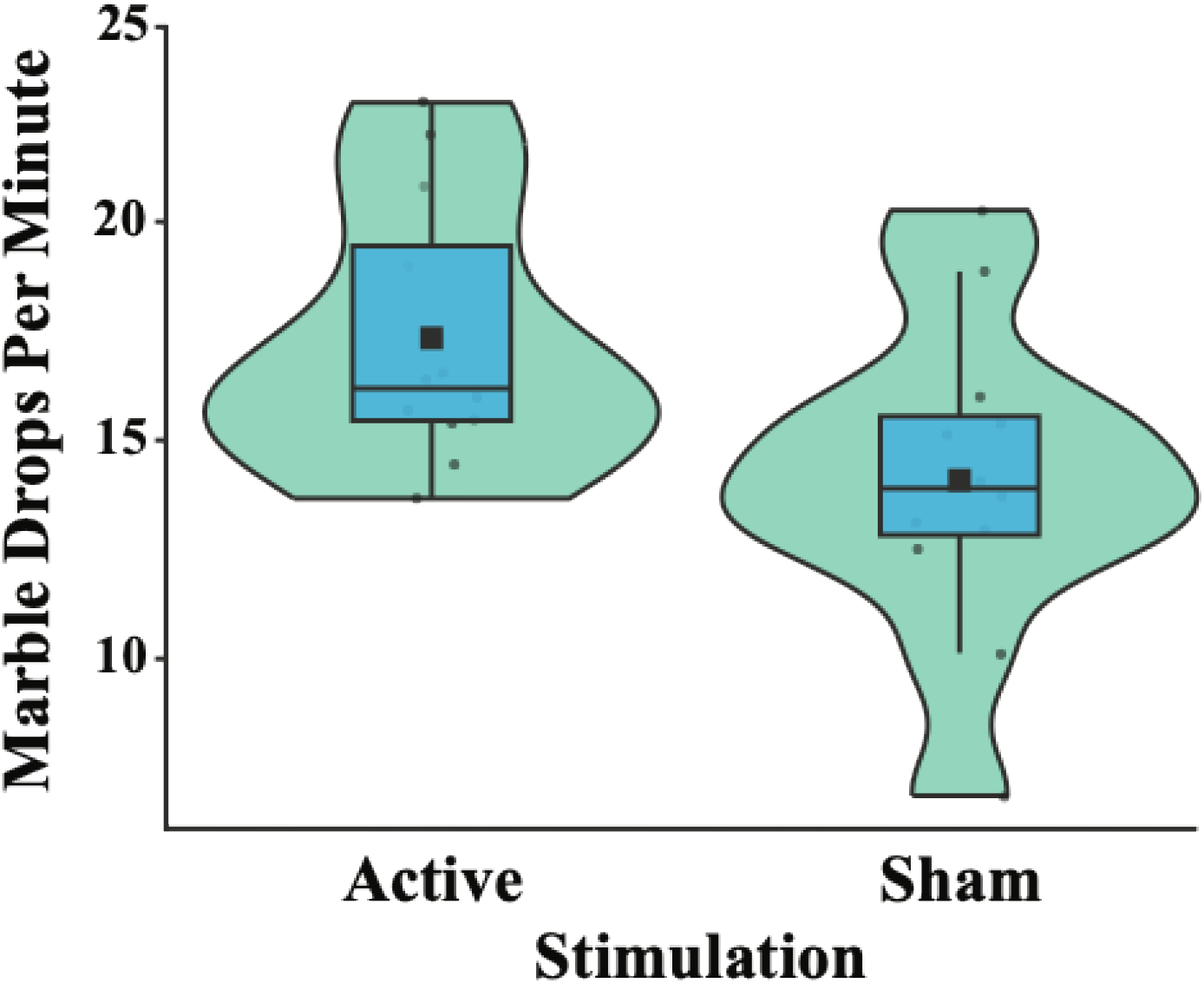
HD-tDCS over the left aSMG enhanced tool use motor learning. Participants receiving active stimulation demonstrate greater performance (marble drops per minute) than those receiving sham stimulation (t(22) = 2.41; p = 0.025), n=12 active, 12 sham, stimulation: ∼3mA active.

**Figure 5.**
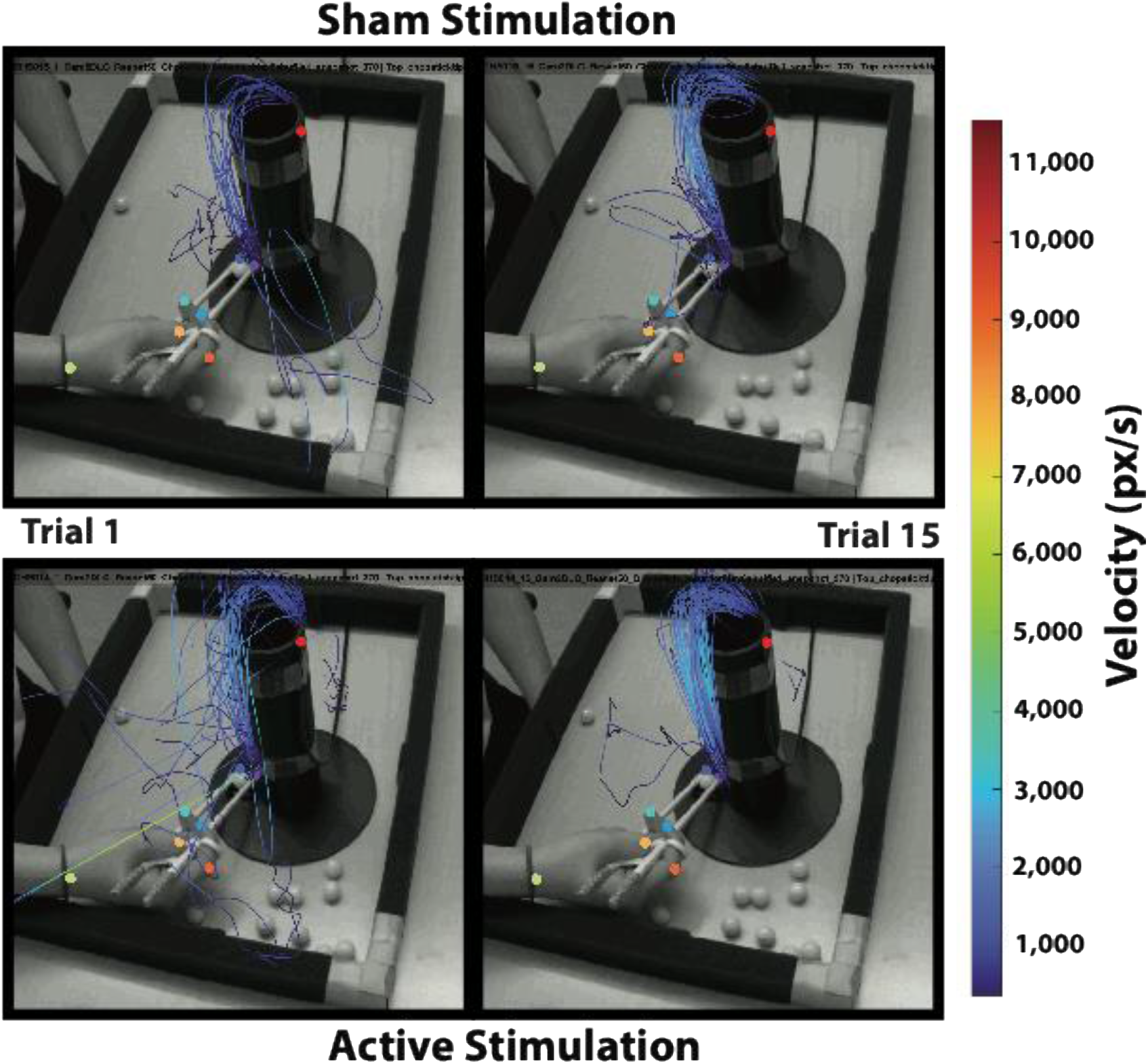
Kinematics measured by movement jerk, total path length, and movement velocity were plotted over still images during performance.

**Figure 6.**
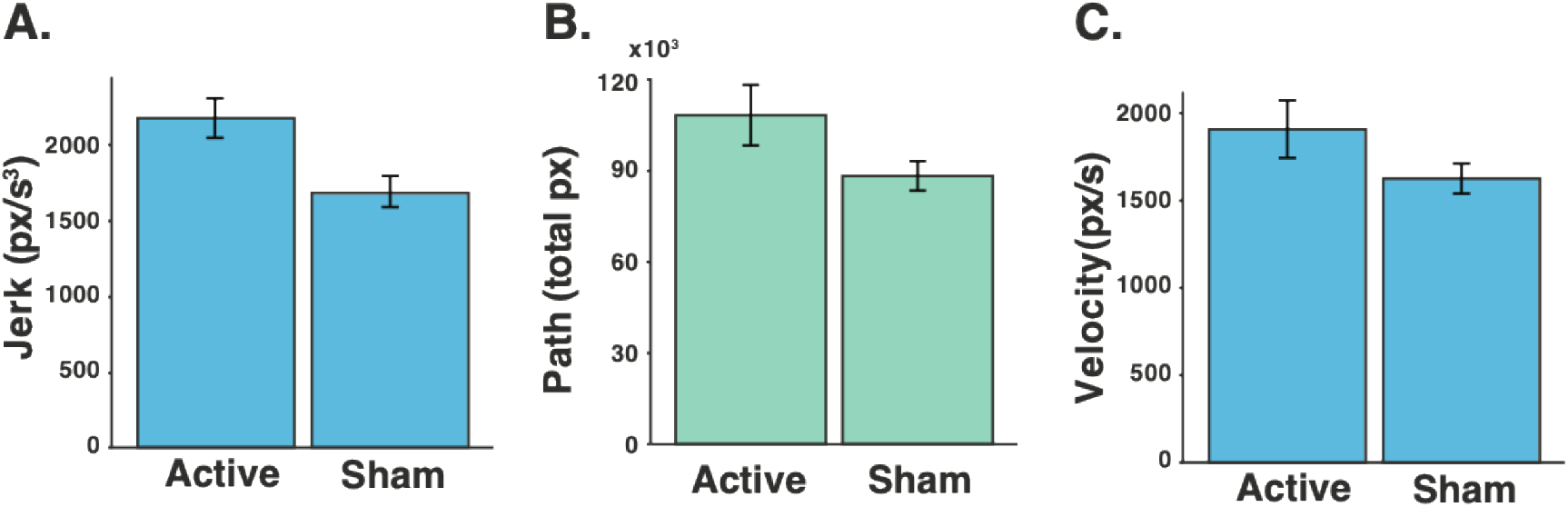
Participants receiving active stimulation had (A) increased movement jerk (21719 vs 16926; t(20.4) = 2.819, p = .010) compared to those receiving sham stimulation. Neither (B) path length (10284 pixels vs. 88299 pixels; t(16.5) = 1.52; p = 0.147) nor (C) movement velocity (1909 pps vs. 1626 pps; t(15.8) = 1.8; p = 0.073) were significantly different between active and sham stimulation. n=12 active, 11 sham.

**Figure 7.**
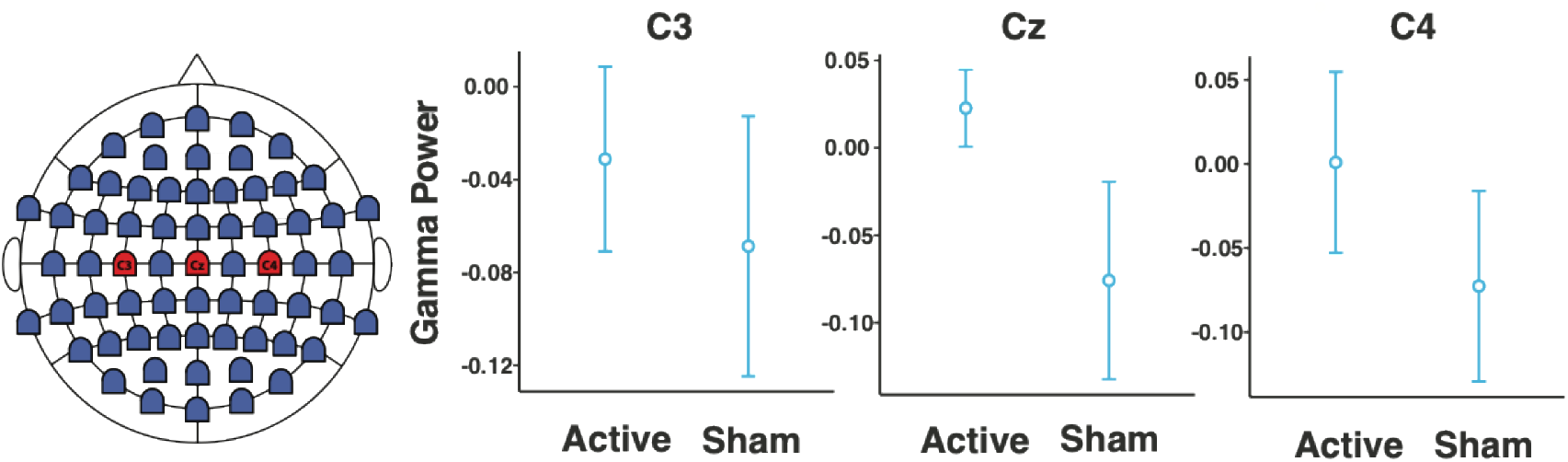
Condition-related differences in task-related gamma-band power were observed over Cz, with higher gamma power during active stimulation (.0227) compared with to sham stimulation (- .0758), (t(14.2) = 3.58, p = .003). No significant differences were observed at C3 (p >.10) or at C4 (t(21.9) = 2.07, p = .051). n=12 active, 12 sham.

## Discussion

The primary purpose of this study was to determine whether modulating activity in the left anterior supramarginal gyrus (aSMG) using HD-tDCS during action observation would enhance the rate of motor learning for a previously unfamiliar tool. Consistent with this hypothesis, both active- and sham-stimulation groups demonstrated significant improvements in MDPM from the first to the final trial, reflecting learning-related gains with repeated practice. However, participants receiving 20 minutes of active HD-tDCS over the aSMG while watching a training-related video exhibited a greater overall improvement across trials relative to the sham group, indicating that aSMG stimulation selectively accelerated the learning process beyond practice effects alone. These findings provide causal evidence supporting the role of the aSMG in facilitating experience-independent tool learning [7, 9] and extend prior work in novel tool use acquisition [27, 28].

The enhanced learning observed in the active stimulation group is consistent with theoretical accounts proposing that the aSMG serves as a key integrative hub within the human tool-use network [7]. Prior work suggests that the aSMG integrates sensorimotor input from the anterior intraparietal sulcus with semantic and technical reasoning processes to support affordance selection and action planning, particularly when encountering unfamiliar tools [7, 29]. By increasing cortical excitability in this region during action observation, HD-tDCS may have strengthened the formation of abstract, transferable representations of tool mechanics prior to direct motor engagement. This mechanism aligns with models of observational learning and associative plasticity, in which neural representations established during observation facilitate subsequent motor execution and learning efficiency.

The kinematic data provide further insight into the nature of the learning enhancements observed in the present study. Participants in the active stimulation group demonstrated an increase in movement jerk, rather than a simple increase in movement velocity. Increased jerk measurements reflect more frequent or larger changes in acceleration. In the context of early and intermediate stages of motor learning, particularly during complex, tool-mediated actions, such increases are often interpreted as reflecting adaptive restructuring of motor commands rather than a degraded smoothness of motor movements. Specifically, heightened jerk can index a transition toward more segmented, information-rich motor control, characterized by sharper initiation, termination, and adjustment phases as internal models are refined. Such kinematic refinements are indicators of motor skill acquisition and are associated with more efficient motor planning, enhanced feedforward control, and reduced reliance on slower, continuous online error correction [30-32].

The electrophysiological findings provide supporting neural evidence that aSMG-targeted HD-tDCS modulated cortical dynamics during the subsequent chopstick task. Specifically, we observed significantly greater task-related gamma-band power at Cz when compared to the sham stimulation protocol. Gamma-band activity over central scalp regions has consistently been linked to movement execution, precision grip control, and the integration of ongoing sensory feedback during skilled action [19-21]. In the context of the chopstick task performed by participants, continuous adjustment of grip force, inter-digit coordination, and online error correction is required, with the elevated central gamma activity likely reflecting more efficient or more engaged local processing to support fine motor control. Importantly, the focal nature of this effect suggests modulation of midline sensorimotor networks, rather than an overall increase in cortical excitability. The absence of a robust beta modulation may indicate that neurostimulation of the aSMG preferentially influenced execution related feedback driven processes, rather than altering broader motor set or inhibitory control mechanisms typically evidenced by changes in beta rhythm. Importantly, this dissociation is in alignment with recent work suggesting gamma oscillations are particularly sensitive to task demands involving fine motor precision and rapid sensorimotor integration whereas beta dynamics may be more closely tied to learning stage, task predictability, or post-movement evaluation [17, 18].

Taken together, these results support the interpretation that aSMG stimulation enhanced motor learning by modulating sensorimotor cortical processing during task performance. When considered alongside the behavioral and kinematic data reported, these findings are consistent with models in which the aSMG contributes to higher-order presentations of tool-related actions, and that once established are used to facilitate more effective sensorimotor integration during execution.

Further study is warranted to explore the broader use and adaptability of HD-tDCS on skill acquisition and rehabilitation, as well as to examine the efficacy of stimulating other brain regions on motor learning, and to characterize the contributions of the aSMG on skill acquisition. Importantly, future studies combining source-resolved electrophysiology or connectivity analyses will be important for clarifying how aSMG-driven modulation interacts with primary and premotor cortices to support skilled tool use.

Some limitations of the present study should be considered when interpreting the findings. First, this study focused on short-term learning within a single experimental session. Although increased marble drops per minute, movement speed, and changes in gamma activity suggest meaningful motor adaptation, the durability of these effects remains unknown. It will be important to determine whether stimulation-related benefits persist over longer retention intervals and whether they generalize to untrained tools or tasks. Longitudinal designs with follow-up assessments are necessary to establish whether aSMG-targeted stimulation produces lasting changes in motor skill acquisition.

Second, given the susceptibility of high-frequency EEG activity to broadband myogenics during hand movements, gamma changes may be reflective of changes in muscular tension and motor noise that would accompany changes in improved dexterity. The central predominance of the gamma reduction suggests that neurostimulation primarily modulated medial sensorimotor and integrative motor control processes rather than producing a uniform suppression across bilateral primary motor cortices. Additionally, the absence of a significant effect at C3 further supports a model in which improved performance reflects altered motor coordination and efficiency, rather than global changes in corticospinal drive or unfiltered myogenic artifact contaminating the EEG signal.

Lastly, although chopstick use provides a well-validated model of novel tool learning, it represents a specific class of fine motor, distal tool use. The extent to which these findings generalize to other categories of tools remains to be determined. Additionally, individual differences in baseline motor proficiency, prior observational experience, or responsiveness to neuromodulation may have contributed to variability in learning rates and should be explicitly examined in future work.

The presented findings demonstrate that HD-tDCS targeting the left anterior supramarginal gyrus during action observation enhances motor learning of a novel tool, chopsticks. Relative to the sham stimulation condition, active stimulation accelerated performance gains, was accompanied by changes in movement execution in alignment with greater efficiency, and also produced focal modulation of gamma-band activity during task performance. Together, this study provides converging behavioral, kinematic, and electrophysiological evidence that the aSMG plays a critical role in supporting skilled tool use. By facilitating the translations of observed actions into effective motor execution, aSMG-targeted stimulation may shorten the time required to learn complex motor skills, including tool use. These findings highlight the potential utility of combining observational learning paradigms with targeted neuromodulation to enhance motor skill acquisition and suggest promising avenues for future applications in both neurorehabilitation and skill training.

## Funding Source

N/A

## Contributors

JS, TB, and LB, contributed to the conception and design of the study. JS, TB, and LB, performed experiments, including data and sample collection. JS, TB, and LB performed data cleaning and analysis. JS organized the datasets. JS, TB and LB performed data cleaning and preprocessing. JS, TB, and LB performed the statistical analysis, and interpreted results of experiments. JS, TB, and LB prepared figures. JS wrote the first draft of the manuscript. JS, TB, and LB wrote sections of the manuscript. All authors contributed to the manuscript revision, read, and approved the submitted version.

## Conflict of Interest

The authors declare that they have no conflict of interest.

## Acknowledgment

The authors would like to recognize the assistance of Paige Pollreisz, Rosara Reicks, and Kamila Haliru in setting up and running the study, as well as the individuals and volunteers who participated in this research.

## Data Availability

The datasets generated during and/or analyzed during the current study are available from the corresponding author upon reasonable request.

## Notes

### Competing Interest Statement

The authors have declared no competing interest.

